# Control of the Endo-Lysosome Homeostasis by the Paracaspase MALT1 regulates Glioma Cell Survival

**DOI:** 10.1101/582221

**Authors:** Kathryn A. Jacobs, Gwennan André-Grégoire, Clément Maghe, Ying Li, An Thys, Elizabeth Harford-Wright, Kilian Trillet, Tiphaine Douanne, Jean-Sébastien Frénel, Nicolas Bidère, Julie Gavard

## Abstract

Glioblastoma is one of the most lethal forms of adult cancer with a median survival of around 15 months. A potential treatment strategy involves targeting glioblastoma stem-like cells (GSC), which constitute a cell autonomous reservoir of aberrant cells able to initiate, maintain, and repopulate the tumor mass. Here, we report that the expression of the paracaspase mucosa-associated lymphoid tissue l (MALT1), a protease previously linked to antigen receptor-mediated NF-κB activation and B-cell lymphoma survival, inversely correlates with patient probability of survival. The knockdown of *MALT1* largely impaired the expansion of patient-derived stem-like cells *in vitro*, and this could be recapitulated with pharmacological inhibitors, *in vitro* and *in vivo*. Blocking MALT1 protease activity increases the endo-lysosome abundance, impaired autophagic flux, and culminates in lysosomal-mediated death, concomitantly with mTOR inactivation and dispersion from lysosomes. These findings place MALT1 as a new druggable target involved in glioblastoma and unveil ways to modulate the homeostasis of endo-lysosomes.

## Introduction

Glioblastoma Multiforme (GBM) represents the most lethal adult primary brain tumors, with a median survival time of 15 months following diagnosis^1,2^. The current standard of care for the treatment of GBM includes a surgical resection of the tumor followed by treatment with alkylating agent temozolomide and radiation. While these standardized strategies have proved beneficial, they remain essentially palliative^1,3,4^. Within these highly heterogeneous tumors exists a subpopulation of tumor cells named Glioblastoma Stem-like Cells (GSCs). Although the molecular and functional definition of GSCs is still a matter of debate, there is compelling evidence that these cells can promote resistance to conventional therapies, invasion into normal brain, and angiogenesis^5,6,7,8,9^. As such, they are suspected to play a role in tumor initiation and progression, as well as recurrence and therapeutic resistance. Owing to their quiescent nature, GSCs resist to both chemotherapy and radiation, which target highly proliferative cancer cells^6,7^. Hence, there is a clear need to identify novel therapeutic targets, designed to eradicate GSCs, in order to improve patient outcome.

GSCs constantly integrate external maintenance cues from their microenvironment, and as such represent the most adaptive and resilient proportion of cells within the tumor mass^5,10^. Niches provide exclusive habitat where stem cells propagate continuously in an undifferentiated state through self-renewal^5^. GSCs are dispersed within tumors and methodically enriched in perivascular and hypoxic zones^11,12,13^. GSCs essentially received positive signals from endothelial cells and pericytes, such as ligand/receptor triggers of stemness pathways and adhesion components of the extracellular matrix^13,14,15,16,17^. They are also protected in rather unfavorable conditions where their stemness traits resist hypoxic stress, acidification, and nutrient deprivation^11,12,18^. Recently, it has been suggested that this latter capacity is linked to the function of the RNA binding protein QKI in the down-regulation of endocytosis, receptor trafficking and endo-lysosome-mediated degradation^18^. GSCs therefore down-regulate lysosomes as one adaptive mechanism to cope with the hostile tumor environment^18^.

Lysosomes operate as central hubs for macromolecule trafficking, degradation, and metabolism^19^. Cancer cells usually show significant changes in lysosome morphology and composition, with reported enhancement in volume, protease activity, and membrane leakiness^20^. These modifications can paradoxically serve tumor progression and drug resistance, while providing an opportunity for cancer therapies. The destabilization of the integrity of these organelles might indeed ignite a less common form of cell death, named as lysosomal membrane permeabilization (LMP). LMP occurs when lysosomal proteases leak into the cytosol and induce features of necrosis or apoptosis, depending on the degree of permeabilization^19^. Recent reports also highlighted that lysosomal homeostasis is essential in cancer stem cell survival^18,21^. Additionally, it has been shown that targeting the autophagic machinery is an effective treatment against apoptosis-resistant GBM^22,23^. The autophagic flux inhibitor chloroquine can decrease cell viability and acts as an adjuvant for TMZ treatment in GBM. However, this treatment might cause neural degeneration at the high doses required for GBM treatment^24^. Therefore, it is preferable to find alternative drugs that elicit anti-tumor responses without the harmful effects on healthy brain cells.

Here, we repurpose several members of a family of drugs, phenothiazines, to disrupt GSC lysosomal homeostasis, induce autophagic features, and subsequently reduce cell survival. Phenothiazines compose a class of neuroleptic and antihistaminic drugs, among which is Mepazine, which has been shown to block the MALT1 cysteine protease^25^. We further established that MALT1 sequesters QKI and maintains low levels of lysosomes, while its inhibition unleashes QKI and hazardously increases endo-lysosomes, which subsequently impairs autophagic flux. This leads to cell death concomitant with mTOR inhibition and dispersion from lysosomes. Disrupting lysosomal homeostasis therefore represents an interesting therapeutic strategy against GSCs.

## Results

### MALT1 Expression sustains Glioblastoma Cell Growth

Glioblastoma Stem-like Cells (GSCs) are suspected to be able to survive outside the protective vascular niche, in a non-favorable environment, under limited access to growth factors and nutrients^11,13^. Because the transcription factor NF-κB is instrumental in many cancers and because it centralizes the paracrine action of cytokines^26,27,28,29^, we revisited The Cancer Genome Atlas (TCGA) for known mediators of the NF-κB pathway that could be aberrantly engaged in GSC expansion. We found that the paracaspase mucosa-associated lymphoid tissue l (MALT1) expression was more significantly correlated with survival than other genes of the pathway (Fig. 1a). This arginine-specific protease is crucial for antigen receptor-mediated NF-κB activation and B-cell lymphoma survival^30,31^. In addition, when GBM patients were grouped by low or high *MALT1* expression levels, there was a significant survival advantage for patients with lower *MALT1* expression (Fig. 1b). Moreover, levels of *MALT1* RNA are elevated in GBM (Grade IV) when compared with lower grade brain tumors (Grades II and III) or non-tumor samples (Fig. 1c-d). At the protein level, MALT1 expression was rather homogenously detectable in four GBM patient-derived cells with stem-like properties (GSCs) from various origins, 2 males (#1 and #12) and 2 females (#4 and #9), mesenchymal (#1 and #4), classical (#9), and neural (#12), and varied in ages from 59 (#12) to 68 (#1 and #9) and 76 years old (#4). MALT1 binding partner BCL10 was also expressed in GSCs, although its level of expression was not correlated with patient probability of survival (Fig. 1a, 1e). Of note, GSCs did not show clear signs of NF-κB activation, as visualized by phosphorylation and degradation of the NF-ΚB inhibitor I Κ B α, as opposed to a treatment with TNFα (Supplementary Fig. 1a). In addition to its scaffold function in the modulation of the NF-κB pathway, MALT1 also acts as a protease for a limited number of substrates^32^. One of MALT1 known substrate, the deubiquitinating enzyme CYLD^33^, was constitutively cleaved in GSCs. This was not further increased upon stimulation with PMA/ionomycin, in contrast to Jurkat T cells, most likely due to a failure to co-opt canonical signaling in this cellular context (Supplementary Fig. 1a). This processing of CYLD was reduced upon treatment with the MALT1 enzymatic inhibitor Mepazine (MPZ)^25^, the competitive inhibitor zVRPR.fmk, or MALT1 siRNA treatment (Supplementary Fig. 1b-d), reinforcing the hypothesis that a fraction of MALT1 is most likely active in growing GSCs, outside its canonical role in antigen receptor signaling and immune cancer cells.

**Fig. 1.**
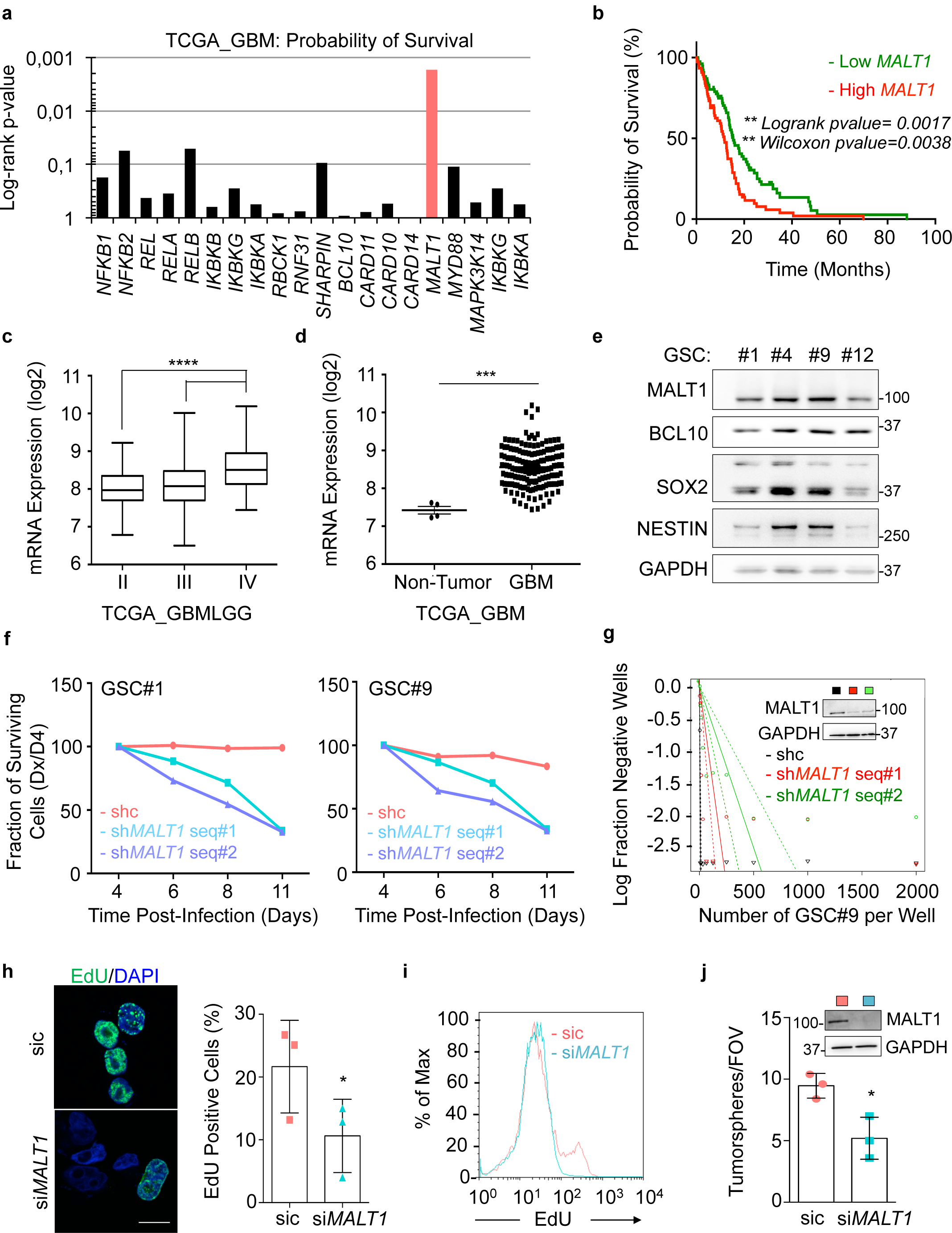
MALT1 Expression sustains Glioblastoma Cell Growth. (a) The Cancer Genome Atlas (TCGA RNAseq dataset) was used on the GlioVis platform^53^ to analyze the probability of survival (log-rank p-value) of 155 GBM patients, for each gene encoding for the well-known mediators of the NF-κB pathway. (b) Kaplan-Meier curve of the probability of survival for patients with low or high *MALT1* RNA level, using median cut-off, based on the TCGA RNAseq dataset. (c-d) Box and whisker plot of *MALT1* mRNA expression in low-grade glioma (grades II and III) or in GBM (grade IV) (TCGA GBMLGG, RNAseq dataset). Alternatively, *MALT1* mRNA expression was plotted in non-tumor samples versus GBM samples (TCGA RNAseq dataset). Each dot represents one human sample. (e) MALT1 and BCL10 protein expression was assessed by western-blot in patient-derived cells with stem-like properties (GSCs, #1 male 68yrs mesenchymal, #4 female 76 yrs mesenchymal, #9 female 68 yrs classical, and #12 male 59 yrs neural). NESTIN and SOX2 stemness markers were also tested. GAPDH served as a loading control. (f) Fraction of surviving cells over time in GSC#1 and GSC#9, transduced with control (shc) or bicistronic GFP plasmids using two different short hairpin RNA (sh*MALT1* sequences, seq #1 and #2). Data are plotted as the percentage of GFP-positive cells at the day of the analysis (Dx), normalized to the starting point (Day 4 post-infection, D4). (g) Linear regression plot of *in vitro* limiting dilution assay (LDA) for control (shc) or sh*MALT1* seq#1 and seq#2 transduced GSC#9. Data are representative of n=2. Knockdown efficiency was verified at day 3 by western-blot using anti-MALT1 antibodies. GAPDH served as a loading control. (h-j) GSC#1 were transfected with non-silencing duplexes (sic) or *MALT1* siRNA duplexes (si*MALT1*) and analyzed 72 hours later. (h) EdU incorporation (green, 2 hours) was visualized by confocal imagery in GSC#1 transfected with sic or si*MALT1* and the percentage of EdU-positive cells was quantified. Nuclei (DAPI) are shown in blue. n> 240 cells per replicate. Scale bar: 10 μm. Data are presented as the mean ± s.e.m. on 3 independent experiments. (i) FACS analysis of EdU staining was performed on similarly treated GSC#9. (j) Tumorspheres per field of view (fov) were manually counted in sic or si*MALT1* transfected GSC#1. Data are presented as the mean ± s.e.m. on 3 independent experiments. Knockdown efficiency was verified at day 3 by western-blot using anti-MALT1 antibodies. GAPDH served as a loading control. All data are representative of n=3, unless specified. Statistics were performed using pairwise comparisons (Tukey’s Honest Significant Difference (HSD) with a 95% confidence interval for panels c and d), and a two-tailed t-test with a 95% confidence interval for panels h and j, *p<0.05, *** p<0.001.

To explore whether MALT1 was engaged in GSCs, the functional impact of MALT1 knockdown was evaluated on proliferation and stemness *in vitro* (Fig. 1f-j). To this end, two individual short hairpin RNA sequences targeting MALT1 (sh*MALT1*) cloned in a lentiviral bi-cistronic GFP expressing plasmid were delivered into GSC#1 and GSC#9 cells. We observed a reduced fraction of GFP-positive cells over time, while cells expressing non-silencing RNA plasmids (shc) maintained a steady proportion of GFP-positive cells, indicating that *MALT1*-silenced cells were proliferating at a lower rate (Fig. 1f). Likewise, cells transfected with si*MALT1* had a lower percentage of EdU-positive cells as compared to non-silenced control cells (Fig. 1h-i). Additionally, GSCs either expressing sh*MALT1* or transfected with si*MALT1* had less stem properties, as evaluated by limited dilution assay and tumorsphere formation (Fig. 1g, 1j). Taken together, these results indicate that MALT1 expression may be important for glioblastoma cell growth.

### Pharmacological Inhibition of MALT1 is Lethal to Glioblastoma Cells

Next, to evaluate the potential of targeting MALT1 pharmacologically, we treated GSCs #1, #4, #9, and #12 with MALT1 inhibitor Mepazine (MPZ)^25^ at a dose of 20 µM. All four GSCs showed a marked reduction in stemness by both limited dilution and tumorsphere assays (Fig. 2a-c). This was accompanied by a marked reduction in the abundance of SOX2 and NESTIN stemness markers (Fig. 2d). Alongside the *in vitro* self-renewal impairment, GSC viability was largely annihilated (Fig. 2e). In contrast, MPZ treatment had no significant effect on viability of brain-originated human cells (endothelial cells, astrocytes, and neurons), ruling out a non-selectively toxic effect (Fig. 2e). In addition, GSC#9 showed an increase in propidium iodide (PI) uptake after MPZ treatment, demonstrating that reduced cell number was due at least partially to cell death (Fig. 2f). Differentiated sister GSCs (DGCs) also showed reduced viability in response to MPZ, indicating that targeting MALT1 may have a pervasive effect on different GBM tumor cells (Fig. 2g). Of note, when *CYLD* and *BCL10* were knocked down through RNA interference, cell viability remained low upon MPZ treatment. This discards an instrumental action of these two MALT1 substrates downstream of MALT1 in MPZ-dependent cell demise (Supplementary Fig. 1e-f).

**Fig. 2.**
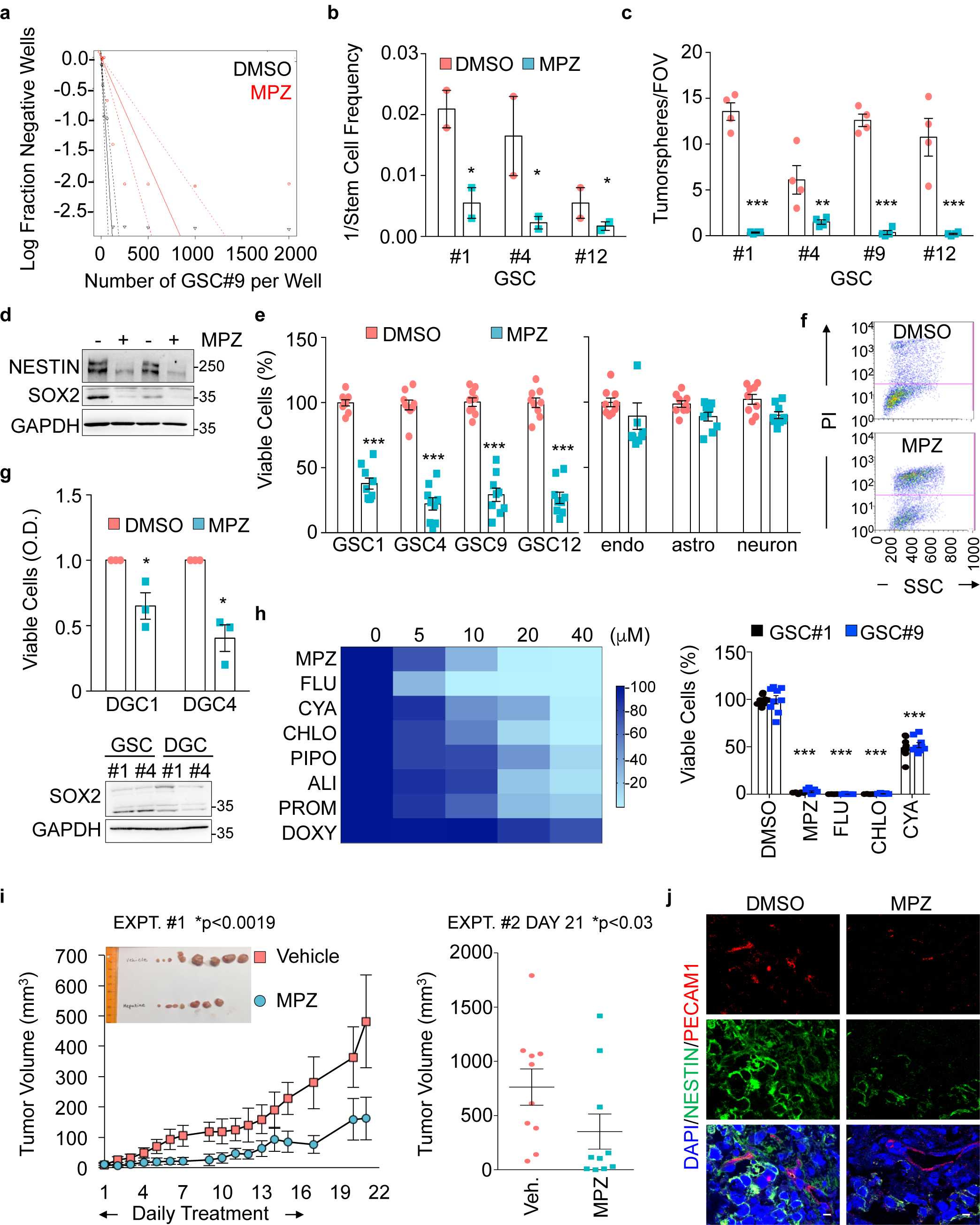
MALT1 Pharmacological Inhibition is Lethal to Glioblastoma Cells. (a) Linear regression plot of *in vitro* limiting dilution assay (LDA) for GSC#9 treated with MALT1 inhibitor, Mepazine (MPZ, 20 μM, 14 days). DMSO vehicle was used as a control. Data are representative of n=2. (b) Stem cell frequency was calculated from LDA in GSCs #1, #4, and #12 treated with MPZ treatment (20 μM, 14 days). Data are presented as the mean ± s.e.m. on 2 independent experiments. (c) Tumorspheres per field of view (fov) were manually counted in GSCs #1, #4, #9, and #12 in response to MPZ (20 μM) and vehicle (DMSO) for 4 days. Data are presented as the mean ± s.e.m. on 4 independent experiments. (d) The expression of the stemness markers SOX2 and NESTIN was evaluated by western-blot in MPZ (+, 20 μM, 16 hours) and vehicle (-, DMSO, 16 hours) treated GSC#9. GAPDH served as a loading control. (e) Cell viability was measured using Uptiblue colormetric assay in GSCs #1, #4, #9, and #12 treated with treatment for 48 hours with vehicle (DMSO) or MPZ (20 μM). Cell viability was measured using Cell TiterGlo luminescent assay in human brain endothelial cells (hCMEC/D3), human astrocytes (SVGp12), and human neurons (SK-N-SH) treated for 48 hours with DMSO or MPZ (20 μM). Data were normalized to their respective DMSO-treated controls and are presented as the mean ± s.e.m of 3 independent experiments in triplicate. (f) FACS analysis of propidium iodide (PI) staining in GSC#9 treated for 48 hours with vehicle (DMSO) or MPZ (20 μM). (g) Cell viability was measured using colormetric MTT assay in differentiated GSC#1 and GSC#4 (DGC) treated for 48 hours with vehicle (DMSO) or MPZ (20 μM). Optical density (O.D.) data were normalized to their respective controls and are presented as the mean ± s.e.m of 3 independent experiments. The expression of the stemness marker SOX2 was evaluated by western-blot in total protein lysates from GSC#1 and GSC#4 and differentiated sister cells DGC#1 and DGC#4. GAPDH served as a loading control. (h) Heatmap of cell viability of GSC#9 using increasing doses (0, 5, 10, 20, 40 μM) of phenothiazines: Fluophenazine (FLU), Cyamemazine (CYA), Chlorpromazine (CHLO), Pipotiazine (PIPO), Alimemazine (ALI), Promethazine (PRO), and Doxylamine (DOXY). Cell viability of GSC#1 and GSC#9 using 20 μM of MPZ, FLU, CHLO, and CYA. Data were normalized to their respective DMSO-treated controls and are presented as the mean ± s.e.m of 3 independent experiments in triplicate. (i) Nude mice were implanted with GSC#9 (10^6^ cells) in each flank and randomized cages were treated with either vehicle (DMSO) or MPZ (8 mg/kg) daily i.p. once tumors were palpable for 14 consecutive days. Tumor volume was measured from the start of treatment until 1 week after treatment was removed. Graph of tumor volume on day 21 post-treatment is presented. Data are presented as the mean ± s.e.m. n=10/group. (j) Cryosections from GSC-xenografted tumors were stained for the endothelial marker PECAM1 (red) and tumor marker NESTIN (green). Nuclei (DAPI) are shown in blue. Scale bar: 20 μm. All data are representative of n=3, unless specified. Statistics were performed using a two-tailed t-test with a 95% confidence interval for panels b to e, a two-way ANOVA with Bonferroni post-test at 95% confidence interval for panel h, and a two-way ANOVA for experiment (Expt) #1 and a Wilcoxon–Mann–Whitney Test for Expt #2 with p-values stated for panel i. *p<0.05 **p<0.01, ***p<0.001.

MPZ is a drug, belonging to the phenothiazine family, and was formerly used in the treatment of schizophrenia^34^. We next evaluated whether other clinically relevant phenothiazines could block MALT1 protease activity and affect GSC viability (Supplementary Fig. 2). Indeed, five of the seven tested phenothiazines reduced MALT1 protease activity upon antigen receptor activation in Jurkat T cells (Supplementary Fig. 2). Furthermore, the effect on MALT1 inhibition was reflected in cell viability, with the best MALT1 protease activity inhibitors (Chlorpromazine and Fluphenazine) having robust effects on cell viability (Fig. 2h). Because MPZ has been shown to efficiently and safely obliterate MALT1 activity in experimental models^30,35^, ectopically implanted GSC#9 mice were challenged with MPZ. Daily MPZ treatment reduced tumor volume in established xenografts, as well as number of NESTIN-positive cells (Figure 2i-j). This effect was prolonged for the week of measurement following treatment withdrawal. Together, these data demonstrate that targeting MALT1 pharmacologically is selectively toxic to GBM cells *in vitro* and *in vivo*.

### MALT1 Inhibition alters Endo-lysosome Homeostasis

To evaluate cell death modality triggered by MALT1 inhibition, caspases activity was blocked with the broad caspase inhibitor Q-VD-OPh (QVD) and increasing doses of MPZ were administered (Supplementary Fig. 3a). QVD treatment did not rescue cell viability upon MPZ treatment, suggesting that cells were not dying through apoptosis, but rather another means. To gain insight into cell death mechanisms, transmission electron microscopy (TEM) was deployed to visualize morphological changes upon MPZ treatment. TEM images showed increased vacuoles and lysosomes compared to control cells (Fig. 3a). The increase in lysosomes could be recapitulated upon si*MALT1* treatment (Supplementary Fig. 3b) In fact, endo-lysosomal protein abundance was amplified upon MALT1 inhibition with MPZ, in a time-dependent manner (Fig. 3b, Supplementary Fig. 3c). Additionally, treatment with the MALT1 competitive inhibitor zVRPR.fmk and other phenothiazines, or *MALT1* knockdown resulted in increased endo-lysosome abundance (Fig. 3c-f, Supplementary Fig. 2c), therefore disqualifying any drug-related action or deleterious accumulation in lysosomes. Conversely, other cellular organelles (early endosomes, mitochondria, golgi, and peroxisomes) remained unchanged upon MPZ treatment (Fig. 3a, Supplementary Fig. 3c-d). These endo-lysosomes appeared to be at least partially functional, as evidenced by pH-based Lysotracker staining, DQ-Ovalbumin and transferrin uptake (Fig. 3d-e, Supplementary Fig. 3e-f). Of note, at a later time point (16 hours) in MPZ treated cells, DQ-Ovalbumin staining was dimmer as compared to early time points (4 hours), which might signify lysosomal membrane permeabilization (Supplementary Fig. 3e). Moreover, ectopic tumors, excised from mice, challenged with a MPZ 2 weeks-regime showed a marked gain in LAMP2 staining and protein amount, as compared to vehicle treated tumors (Fig. 3g). Our data demonstrated that MALT1 knockdown and pharmacological inhibition provoke a meaningful endo-lysosomal increase.

**Fig. 3.**
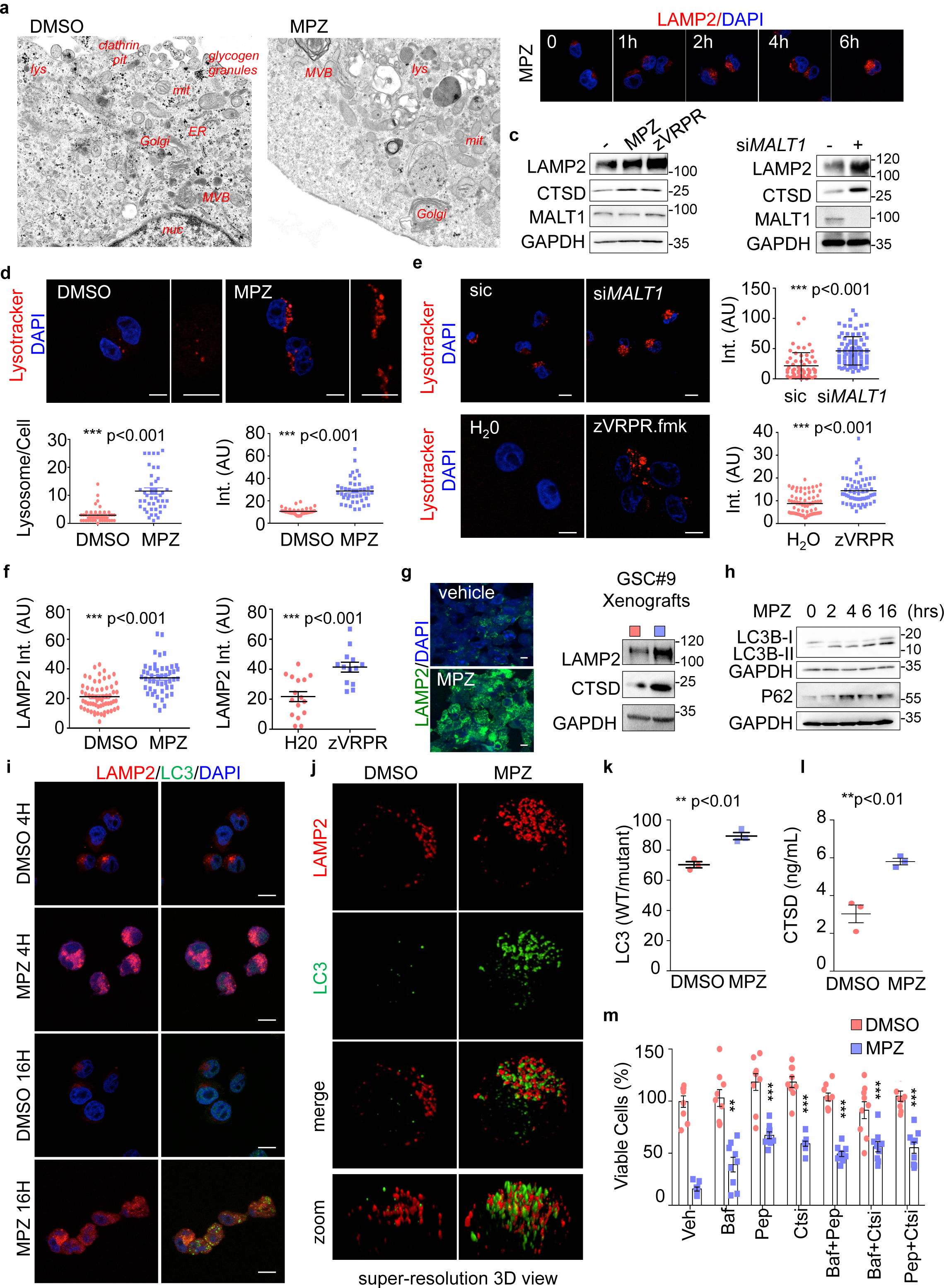
MALT1 Pharmacological Inhibition alters Endo-Lysosome Homeostasis. (a) Transmission electron microscopy of GSC#9 treated with vehicle (DMSO) or MPZ (20 μM) for 16 hours. ER: endoplasmic reticulum; MVB: multivesicular bodies; lys: lysosome; mit: mitochondria; nuc: nucleus. (b) Confocal analysis of LAMP2 staining (red) at 0, 1, 2, 4, and 6 hours post-MPZ (20 μM) treatment. Nuclei (DAPI) are shown in blue. Scale bar: 10 μm. (c) Western-blot analysis of LAMP2, CTSD, and MALT1 in total protein lysates from GSC#9 treated for 16 hours with MPZ (20 μM) or zVRPR.fmk (75 µM). DMSO was used as vehicle. Alternatively, western-blot analysis of LAMP2, MALT1, and CTSD was performed in total protein lysates from GSC#9 transfected with non-silencing duplexes (sic) or *MALT1* targeting siRNA duplexes (si*MALT1*). GAPDH served as a loading control. (d-e) Confocal analysis of Lysotracker staining (red) in GSC#9 treated for 16 hours with vehicle (DMSO) or MPZ (20 μM). Alternatively, GSC#9 were either transfected with sic or si*MALT1* (upper panel) or treated for 16 hours with H_2_O or zVRPR.fmk (75 µM) (bottom panel). Both the number of lysotracker-positive puncta and the lysotracker pixel intensity (Int. in arbitrary unit, A.U.) were quantified per cell. Each dot represents one cell. n>30. Nuclei (DAPI) are shown in blue. Scale bars: 10 μm. (f) Quantification of LAMP2 staining pixel intensity on GSC#9 treated for 16 hours with vehicle (DMSO or H_2_O), MPZ (20 µM) or zVRPR.fmk (75 µM). Each dot represents one cell. n>30. (g) Cryosections from GSC#9-xenografted tumors in vehicle and MPZ-challenged animals (as described in panel 2h) and assessed for LAMP2 staining (green). Nuclei (DAPI) are shown in blue. Scale bar: 10 μm. Western-blot analysis of LAMP2 and CTSD was performed in tumor lysates. GAPDH served as a loading control. (h) Western-blot analysis of LC3B and P62 in total protein lysates from GSC#9 at 0, 2, 4, 6, and 16 hours post-MPZ treatment (20 µM). GAPDH served as a loading control. (i) Confocal analysis of LAMP2 (red) and LC3B (green) in GSC#9 treated for 4 and 16 hours with vehicle (DMSO) and MPZ (20 µM). Nuclei (DAPI) are shown in blue. Scale bars: 10 μm. (j) Super-resolution imaging (SIM, Structured Illumination Microscopy) of LAMP2 (red) and LC3B (green) staining in GSC#9 treated for 16 hours with vehicle (DMSO) or MPZ (20 µM). (k) GSC#9 were transfected with LC3B reporters (wild type WT or G120A mutant which cannot be lipidated), treated 24 hours later with vehicle (DMSO) or MPZ (20 µM) for 6 more hours. Ratios of WT/mutant luciferase signals are presented as the mean ± s.e.m of 3 independent experiments. (l) CSTD ELISA was performed on culture media from GSC#9 treated for 8 hours with vehicle (DMSO) or MPZ (20 µM). Data are presented as the mean ± s.e.m of 3 independent experiments. (m) Cell viability was measured using Cell TiterGlo luminescent assay in GSC#9 treated for 48 hours with vehicle (DMSO) or MPZ (10 μM), following a 30 minute-pre-treatment with the following drugs: Bafilomycin A1 (Baf, 100 nM), Pepstatin A (Pep, 1 μg/mL), or CTS inhibitor 1 (Ctsi, 1 μM). Data were normalized to the vehicle-treated controls and are presented as the mean ± s.e.m of 3 independent experiments in triplicate, stars refer to comparison to Vehicle + MPZ group (blue squares). All data are representative of n=3, unless specified. Statistics were performed using a two-tailed t-test with a 95% confidence interval for all experiments with p-values stated, except panel m, which used a two-way ANOVA with Bonferroni post-test at 95% confidence interval. **p<0.01, ***p<0.001.

Because autophagy is fueled by endo-lysosome activity, we also explored the impact of MALT1 inhibition on autophagy in GSCs, as estimated by LC3B modifications. The turnover of LC3B and the degradation of the autophagy substrate P62 also reflect autophagic flux^36^. Upon MPZ treatment, there was an accumulation of lipidated LC3B (LC3B-II) and P62 protein amount over time, suggesting impaired autophagic flux (Fig. 3h). Treatment with MPZ also led to a significant increase in LC3B puncta at later time points (16 hours), subsequent to lysosomal increase (4 hours) (Fig. 3i). Super-resolution microscopy using structured illumination microscopy (SIM) further revealed that these LC3 structures were covered with LAMP2-positive staining (Fig. 3j). This was concomitant with a reduced LC3B turnover, as evaluated via luciferase assay (Fig. 3k). Our data suggest that lysosomal increase upon MALT1 inhibition precedes autophagic flux impairment. At the functional level, there was increased CTSD release by GSCs treated with MPZ, implying lysosomal membrane permeabilization (Fig. 3l). Accordingly, treatment with lysosomal inhibitors partially rescued cells from MPZ-induced cell death (Fig. 3m). Thus, the protease activity of MALT1 appears to be required to maintain innocuous endo-lysosomes in GSCs.

### MALT1 modulates the Lysosomal mTOR Signaling Pathway

In order to further characterize the mode of action of MALT1 inhibition in GSCs, we performed RNA-sequencing analysis on GSCs treated with MPZ for 4 hours, prior to any functional sign of death. Our results showed 7474 differentially expressed genes, among which no obvious endo-lysosomal protein encoding genes were identified, which was further confirmed by qPCR (Fig. 4a-e, Supplementary Table 1). Of note, VGF, recently shown to promote GSC/DGC survival, was down-regulated upon MPZ treatment^37^ (Fig. 4e, Supplementary Table 1). In line with a non-transcriptional regulation of lysosome biogenesis, knockdown of the master regulator of lysosomal transcription TFEB^38^ failed to reduce autophagy signature and lysosomal protein up-regulation upon MPZ treatment (Fig. 4f). We thus hypothesized that the observed endo-lysosomal increase was due to modulation in their translation and/or RNA metabolism. When translation was blocked with cycloheximide, MPZ failed to increase endo-lysosomal protein amounts (Supplementary Fig. 4a). Likewise, RNAseq analysis unveiled putative changes in translation (peptide chain elongation, ribosome, cotranslational protein targeting, 3’-UTR mediated translational regulation), RNA biology (influenza viral RNA, nonsense mediated decay), metabolism (respiratory electron transport, ATP synthesis, oxidative phosphorylation, respiratory electron transport), and an mTOR signature (Bilanges serum and rapamycin sensitive genes) (Fig. 4a-d). Because mTOR sustains GSC expansion (Supplementary Fig. 4b) and its activation is linked to lysosomal biogenesis^14,39,40^, we further explored this possibility. Notably, MALT1 activity has been shown to participate in mTOR activation upon antigen receptor engagement, although the mechanism of action remains poorly understood^41,42^. In fact, MPZ and phenothiazine pharmacological challenge, as well as *MALT1* siRNA blunted mTOR activation in GSCs, as evaluated through the activation and phosphorylation of AKT, p70S6K and S6 ribosomal protein (Fig. 4g-j). MPZ treatment also reduced inhibitory phosphorylation of autophagy regulator ULK1 at serine 757 (Fig. 4g), which may partially account for increased autophagic features upon MPZ treatment. Furthermore, as phosphorylation of 4EBP1 increases protein translation by releasing it from EIF4E^43^, and as it can be resistant to mTOR inhibition^44^, we evaluated 4EBP1 phosphorylation levels over time in response to MPZ (Supplementary Fig. 4c). Although reduced shortly upon MPZ addition, phosphorylation returned at later time points, which may allow for the observed translational effect despite mTOR inhibition. As mTOR signaling is intimately linked to lysosomes^45^, we explored the impact of MPZ treatment on mTOR positioning. MPZ challenge induced a striking dispersion of mTOR foci (Fig. 4k). In fact, mTOR staining divorced LAMP2-positive structures (Fig. 4l, Supplementary Fig. 4d). Interestingly, TFEB silencing influenced neither lysosome protein intensification nor mTOR recruitment at lysosomes (Fig. 4f, Supplementary Fig. 4d). Conversely, mTOR staining was dispersed from endo-lysosomes upon MPZ, zVRPR.fmk, or phenothiazines treatment, which likely accounts for decreased mTOR activity. These results suggest that MALT1 affects lysosomal homeostasis post-transcriptionally, and that this increase coincides with weak mTOR signaling, which may be due to displacement of mTOR from its lysosomal signaling hub.

**Fig. 4.**
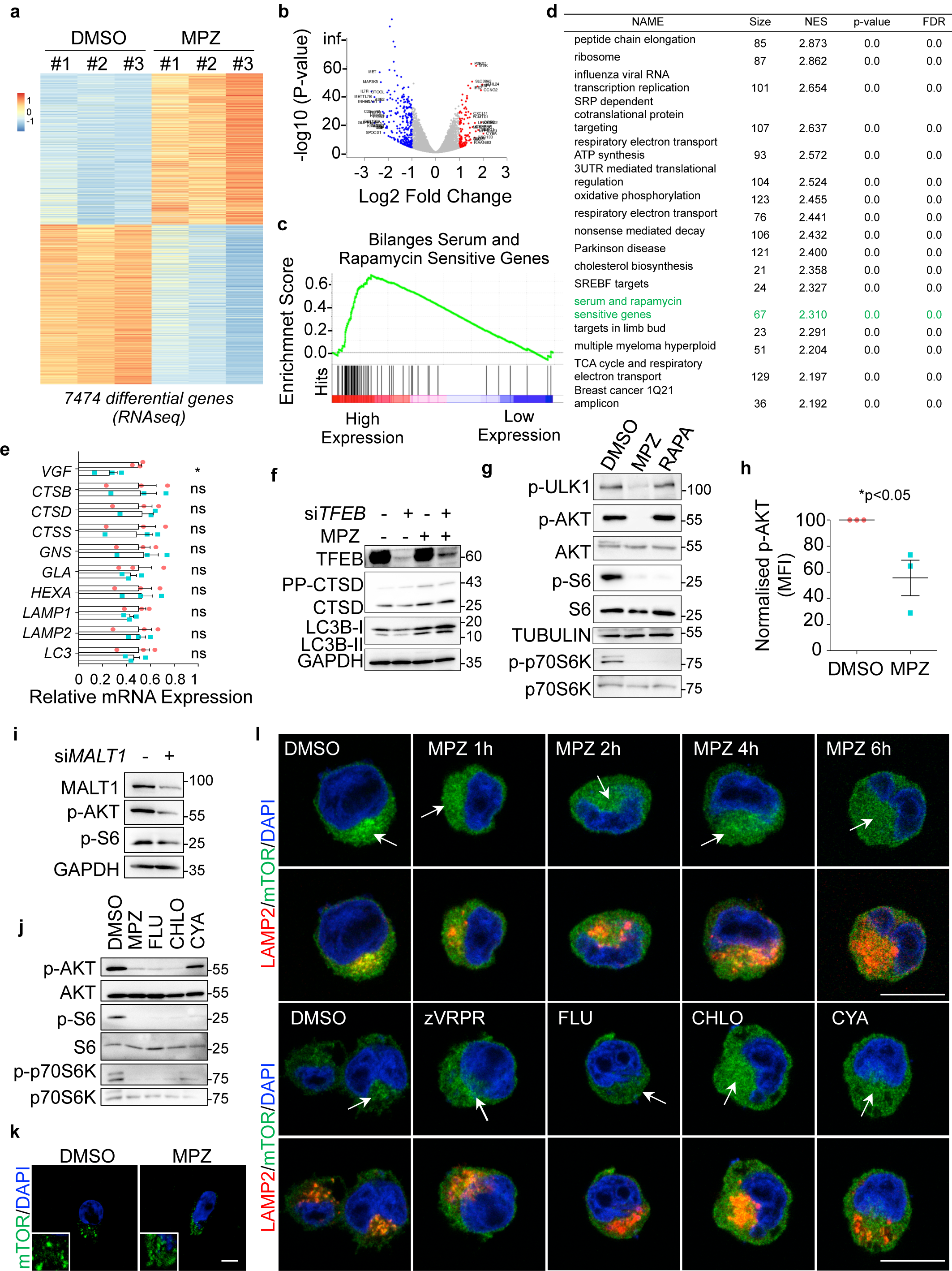
MALT1 modulates the Lysosomal mTOR Signaling Pathway. (a) Heatmap of differentially expressed genes obtained from RNAseq analysis of GSC#9 treated for 4 hours with vehicle (DMSO) or MPZ (20 μM), from three biological replicates. (b) Volcano plot of differentially expressed genes in RNAseq analysis of GSC#9, expressed as fold changes between vehicle (DMSO) and MPZ-treated cells. (c) GSEA (gene set enrichment analysis) plot showing enrichment of “Bilanges serum and rapamycin sensitive genes” signature in vehicle (DMSO) versus MPZ-treated triplicates. (d) Table of top differential pathways in DMSO versus MPZ-treated triplicates. Size of each pathway, normalized enrichment scores (NES), p-value and false discovery rate q value (FDR) were indicated. (e) qRT-PCR was performed on total RNA from GSC#9 treated for 4 hours with vehicle (DMSO) or MPZ (20 μM). Histograms showed changes in RNA expression of indicated targets. Data were normalized to two housekeeping genes (*ACTB, HPRT1*) and are presented as the mean ± s.e.m of technical triplicates. *p<0.05. (f) Western-blot analysis of LC3B, CSTD and TFEB in total protein lysates from GSC#9 transfected with non-silencing duplexes (sic) or siRNA duplexes targeting *TFEB* (si*TFEB*) and treated with vehicle (DMSO) or MPZ (20 μM) for 16 hours. GAPDH served as a loading control. (g) Western-blot analysis of p-ULK1, p-AKT, p-S6, and p-p70S6K in GSC#1 treated for 1 hour with MPZ (20 μM) or Rapamycin (RAPA, 50?nM). Total AKT, S6, p70S6K and GAPDH served as loading controls. DMSO was used as a vehicle. (h) FACS analysis of p-S473AKT in GSC#9 treated for 1 hour with vehicle (DMSO) or MPZ (20 μM). MFI (Mean Fluorescent Intensity) are normalized to vehicle treated controls and are presented as the mean ± s.e.m. on 3 independent experiments. (i) Western-blot analysis of MALT1, p-AKT, and p-S6 in total protein lysates from GSC#9 transfected with non-silencing duplexes (sic) or *MALT1* targeting siRNA duplexes (si*MALT1*). GAPDH served as a loading control. (j) Western-blot analysis of p-AKT, p-S6, and p-p70S6K in total protein lysates from GSC#9 treated for 1 hour with vehicle (DMSO) or 20 μM of phenothiazine compounds (MPZ, FLU, CHLO, and CYA). Total AKT, total S6 and total p70S6K served as loading controls. (k) Confocal analysis of mTOR staining in GSC#9 cells that received either vehicle (DMSO) or MPZ (20 µM, 6 hours). Cells were treated with saponin (0.1%, 10 sec) prior fixation. Nuclei (DAPI) are shown in blue. Scale bars: 10 μm. (l) Confocal analysis of LAMP2 (red) and mTOR (green) staining in GSC#9 treated with vehicle (DMSO) or zVRPR.fmk (75 μM), FLU (20 μM), CHLO (20 μM), and CYA (20 μM). Nuclei (DAPI) are shown in blue. Arrows point to LAMP2-positive area. Scale bars: 10 μm. All data are representative of n=3, unless specified. Statistics were performed using a two-tailed t-test with a 95% confidence interval for all experiments with p-values stated.

### MALT1 is Negatively linked to the Endo-lysosomal Regulator QKI

Shinghu et al.^18^ recently demonstrated that the RNA binding protein Quaking (QKI) regulates endo-lysosomal levels in GBM. They showed that GBM-initiating cells maintain low levels of endo-lysosomal trafficking in order to reduce receptor recycling. QKI was suggested to regulate RNA homeostasis of endo-lysosome elements, independently of the TFEB-driven endo-lysosome biogenesis^18^. As our data suggest a counterbalancing role of MALT1 in lysosomal biogenesis, we revisited the TCGA and compared the expression of *MALT1* with that of *QKI* in GBM patients. Interestingly, there was a negative correlation between the expression of the two genes (Fig. 5a). In addition, *QKI* and *MALT1* were both linked to the expression of 7 common lysosomal lumen genes (Fig. 5a). This prompted us to examine QKI pattern in GBM. First, QKI was indeed expressed in a panel of GSCs, as well as in ectopic xenografts (Supplementary Fig. 4e). As expected, QKI displayed cytosolic and nuclear forms^46^, as evidenced by cellular fractionation and immunofluorescence (Supplementary Fig. 4f-g). Similarly, human GBM samples from 2 patients showed pervasive QKI staining (Supplementary Fig. 4h). Given these findings, we decided to explore the possible link between MALT1 and QKI in GSCs. Co-immunoprecipitation experiments were thus deployed using QKI and the MALT1 binding partner BCL10 as baits. This showed that MALT1 was pulled down with QKI in GSC#1 and GSC#9, and *vice versa* (Fig. 5b). Binding was, however, reduced in cells exposed to MPZ for 1 hour (Fig. 5c). This suggests that active MALT1 tethered QKI in GSCs, while blocking MALT1 unleashed a fraction of QKI from the BCL10/MALT1 complex.

**Fig. 5.**
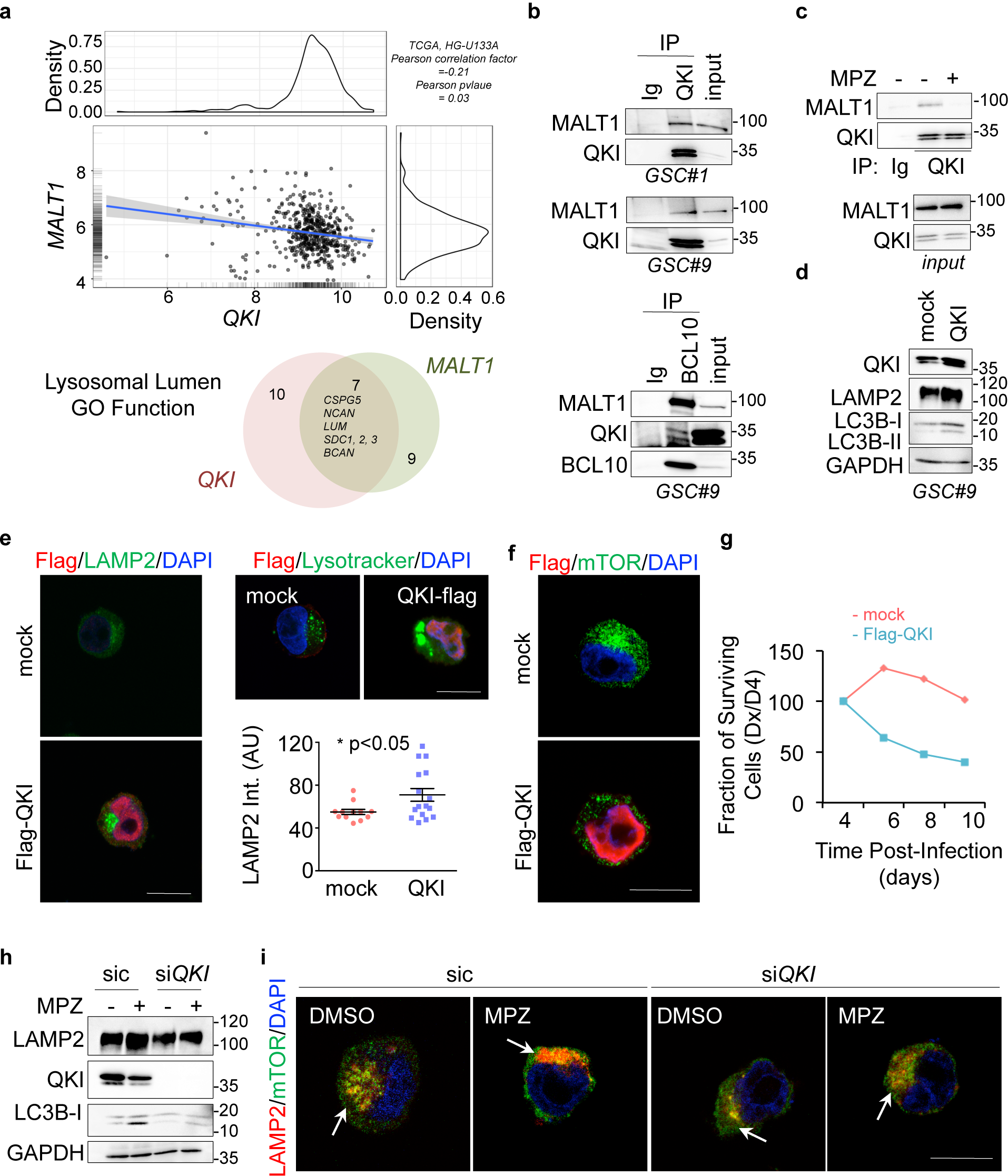
MALT1 is Negatively linked to the Endo-Lysosomal Regulator QKI. (a) Correlation between *MALT1* and *QKI* expression was analyzed using The Cancer Genome Atlas (TCGA, HG-U133A dataset) on the GlioVis platform^53^. Pearson correlation factor=-0.21, p-value=0.03. Differential expression analysis related to either *MALT1* or *QKI* expression highlighted a lysosomal lumen GO function. Venn diagram of overlapping lysosomal enriched protein encoding genes from this comparison showed 7 shared genes, together with 9 and 10 specific genes for *MALT1* and *QKI* expression, respectively. (b) GSC#1 and GSC#9 protein lysates (input) were processed for immunoprecipitation (IP) using control immunoglobulins (Ig), anti-QKI or anti-BCL10 antibodies. Input and IP fractions were separated on SDS-PAGE and western-blots were performed using anti-MALT1, anti-QKI and anti-BCL10 antibodies, as specified. (c) Total protein lysates (input) from GSC#9 treated with vehicle (-, DMSO) or MPZ (+, 20 μM, 1 hour) were processed for control immunoglobulins (Ig) or anti-QKI antibodies immunoprecipitation (IP). Western-blots were performed with indicated antibodies. (d) Western-blot analysis of QKI, LAMP2 and LC3B in GSC#9 overexpressing either empty vector (mock) or Flag-QKI. GAPDH served as a loading control. (e) Confocal analysis of LAMP2 (green) or Flag (red) in GSC#9 overexpressing either empty vector (mock) or Flag-QKI. Alternatively, Lysotracker (green) was shown. Nuclei (DAPI) are shown in blue. Scale bars: 10 μm. Quantification of LAMP2 staining pixel intensity on GSC#9 transfected with mock and Flag-QKI. Each dot represents one cell. n>15. f) Confocal analysis of mTOR (green) or Flag (red) in GSC#9 overexpressing either empty vector (mock) or Flag-QKI. Nuclei (DAPI) are shown in blue. Scale bars: 10 μm. (g) Fraction of surviving cells over time in GSC#1 and GSC#9, transduced with empty vector (mock) or Flag-QKI bicistronic GFP plasmids. Data are plotted as the percentage of GFP-positive cells at the day of the analysis (Dx), normalized to the starting point (Day 4 post-infection, D4). Data are representative of n=3. (h) GSC#9 transfected with non-silencing RNA duplexes (sic) or *QKI* targeting siRNA duplexes (si*QKI*) were treated for 16 hours with vehicle (DMSO) or MPZ (20 μM). Total protein lysates were processed for western-blots against LAMP2, CTSD, QKI, and LC3B expression, as indicated. GAPDH served as a loading control. (i) Confocal analysis of mTOR (green), LAMP2 (red) in GSC#9 transfected with sic or si*QKI*, and treated for 16 hours with vehicle (DMSO) or MPZ (20 μM). Nuclei (DAPI) are shown in blue. Scale bars: 10 μm. All data are representative of n=3, unless specified. Statistics were performed using a two-tailed t-test with a 95% confidence interval for all experiments with p-values stated. *p<0.05.

To next challenge the function of this putative neutralizing interaction of MALT1 on QKI, QKI expression was manipulated to alter QKI/MALT1 stoichiometry in GSCs. Strikingly, transient overexpression of QKI phenocopied the effect of MALT1 inhibition on endo-lysosomes. Reinforcing pioneer findings of QKI action on endo-lysosome components in transformed neural progenitors^18^, ectopically expressed QKI was sufficient to increase LAMP2 protein amount and lipidated LC3B (Fig. 5d-e). Accordingly, the augmented lysosome staining synchronized with mTOR displacement (Fig. 5f). Corroborating the surge of lysosomes, the fraction of cells overexpressing QKI was drastically reduced over time, while fraction of cells expressing an empty vector remained stable, suggesting that exacerbated QKI expression hampered cell proliferation (Fig. 5g). Conversely, cells knocked down for *QKI* did not show the same MPZ-driven increase in LAMP2 and lipidated LC3B (LC3B-II), suggesting that *QKI* knockdown can partially rescue cells from endo-lysosomal increase (Fig. 5h-i). Reinforcing this idea, mTOR staining was no longer dissipated from lysosomes upon MPZ treatment without QKI (Fig. 5i). Thus, *QKI* silencing rescued endo-lysosomes equilibrium in MPZ-treated cells, further indicating that MALT1 is negatively linked to the endo-lysosomal regulator QKI.

## Discussion

Here we provide evidence that the paracaspase MALT1 activity is decisive for growth and survival in GBM. Our data indicate that MALT1 inhibition causes indiscipline of endo-lysosomal and autophagic proteins, which appears to occur in conjunction with a deficit in mTOR activity. In addition to the known MALT1 inhibitor Mepazine^25^, we show that several other clinically relevant phenothiazines can potently suppress MALT1 enzymatic activity, and have similar effects to MPZ on endo-lysosomes and cell death in GSCs. Since these drugs efficiently cross the blood-brain barrier in humans^47^ and since they are currently used in the clinic, they represent an exciting opportunity for drug repurposing.

The disruption of endo-lysosomal homeostasis appears to be the cause of death upon MALT1 inhibition in GSCs. As CSTD release is accelerated upon MALT1 blockade, and as inhibitors of lysosomal cathepsins (cathepsin inhibitor 1 and pepstatin A), but not pan-caspase blockade (QVD), can partially rescue cell viability, we hypothesize that cells may be dying from a form of caspase-independent lysosomal cell death (LCD)^19^. During this form of death, which may also be initiated by cathepins, lysosomal membrane permeabilization (LMP) allows cathepsins to act as downstream mediators of cell death upon leakage into the cytosol^19^. Additional studies will determine how exactly MALT1 inhibition drives lysosomal death in GSCs. Nevertheless, we found that inhibition of cathepsins provides only partial protection to cells treated with MPZ (Fig. 3m). Autophagic features may also play a part in cell death. Induction of autophagy likely occurs due to reduced inhibition of ULK1 (Fig. 4g) as a consequence of mTOR dispersion from lysosomes^39,40^ (Fig. 4l). Whether inducing or blocking autophagy are preferable therapeutic strategies in treating GBM remains hotly debated, with some groups reporting beneficial effects of blocking autophagy, and others preferring its activation as a therapeutic strategy^48,23^. Here we show that the observed increased autophagic features are associated with reduced autophagic flux. Impairment in autophagic flux reduces a cell’s ability for bulk degradation^36^. Others have shown that lysosomal dysfunction, such as LMP, can impede upon autophagic flux and eventually lead to cell death^49,50^. Because of this, we infer that reduced autophagic flux is a downstream consequence of LMP, and ultimately contributes to LCD in our cells.

MALT1 has previously been linked to mTOR activity^41,42^. For instance, MALT1 was reported to be necessary for glutamine uptake and mTOR activation after T cell receptor engagement^41^. As a consequence, the inhibition of MALT1 with zVRPR.fmk causes a reduction in the phosphorylation of S6 and p70S6K^42^. Our data now extend these findings to GSCs, however, the exact mechanism by which this occurs remains to be explored in both cellular backgrounds. Immunofluorescence analysis of mTOR positioning after MPZ treatment suggests that inhibition of mTOR is linked to its dispersion from the lysosomes, concurrent with lysosomal increase. However, we and others speculate that there may exist unidentified substrates of MALT1, which link its protease activity directly to mTOR activation^32^. This may also rationalize the need for constitutive MALT1 activity in GSCs, as mTOR is constantly active in these cells^14^. One hypothesis is that mTOR inhibition and/or dissociation from lysosomes originate from lack of processing of unknown MALT1 substrates and is then exacerbated once homeostasis is disrupted.

How is QKI involved? Based on our data, we hypothesize that MALT1 sequesters QKI to prevent it from carrying out its RNA-binding function. Interestingly, MALT1 is already known to regulate other RNA-binding proteins Regnase-1/ZC3H12A, Roquin-1/RC3H1 and Roquin-2/RC3H2^51,52^. We propose that upon MALT1 inhibition QKI is released and free to bind its RNA binding partners. QKI has already been shown to bind directly to lysosomal RNAs in progenitor cells^18^. It is thus tempting to speculate that QKI-dependent stabilization of lysosomal RNAs would preference the system towards more translation of these genes upon MALT1 inhibition. Indeed our RNA sequencing data suggests changes in translation and RNA biology upon MPZ treatment, however further study is needed to validate whether there is increased QKI binding to lysosomal RNAs upon MALT1 inhibition. Notably, QKI dependent lysosomal increase appears to be a post-transcriptional effect, independent of TFEB. As such, we propose a method of dual lysosomal control in GSCs whereby transcriptional biogenesis is tightly checked by known mTOR/TFEB pathway, and MALT1 acts on post-transcriptional regulation by isolating QKI from RNAs (Supplementary Fig. 4i).

These findings place MALT1 as a new druggable target operating in non-immune cancer cells and involved in endo-lysosome homeostasis. Lysosomal homeostasis appears vital for glioblastoma cell survival and thus presents an intriguing axis for new therapeutic strategies in GBM.

## Materials and Methods

### Ethics statement

Informed consent was obtained from all patients prior to sample collection for diagnostic purposes. This study was reviewed and approved by the institutional review boards of Sainte Anne Hospital, Paris, France, and Laennec Hospital, Nantes, France, and performed in accordance with the Helsinki Protocol. Animal procedures were conducted as outlined by the European Convention for the Protection of Vertebrate Animals used for Experimental and other Scientific Purposes (ETS 123) and approved by the French Government (APAFIS#2016-2015092917067009).

### The Cancer Genome Atlas (TCGA) Analysis

The Cancer Genome Atlas (TCGA) was explored via the Gliovis platform (http://gliovis.bioinfo.cnio.es/)^53^. RNAseq databases (155 patients) were used to interrogate data related to *MALT1* expression (levels of RNA, probability of survival, correlation with *QKI* expression). Optimal cut-offs were set. All subtypes were included and histology was the only selective criteria.

### Cell Culture, siRNA and DNA transfection, lentiviral transduction

GBM patient-derived cells with stem-like properties (GSCs) were isolated as previously described^15,54^. Mesenchymal GSC#1 and #4, Classical GSC#9, and Pro-Neural GSC#12 were cultured as spheroids in NS34 medium (DMEM-F12, with N2, G5 and B27 supplements, glutamax and antibiotics, Life Technologies). In order to induce differentiation, GSCs were grown in DMEM with 10% fetal bovine serum (FBS), glutamax and antibiotics (Life Technologies), for at least 2 weeks. Differentiation of sister cells (DGC) were monitored through their morphology and NESTIN and/or SOX2 loss of expression. Human brain microvascular endothelial cells (hCMEC/D3, a gift from PO Couraud, Institut Cochin, Paris, France) and HEK-293T (ATCC, LGC Standards) were cultured as previously described^54^. SVG-p12 and SK-N-SH (ATCC, LGC Standards) were cultured in MEM with 10% fetal bovine serum (FBS), and antibiotics (Life Technologies).

Stealth non-silencing control duplexes (low-GC 12935111, Life Technologies), and small interfering RNA duplexes (Stealth RNAi, Life Technologies) were transfected using RNAiMAX lipofectamine (Life Technologies). The following duplexes targeting the respective human genes were: CAGCAUUCUGGAUUGGCAAAUGGAA (*MALT1*), CCTTGAGTATCCTATTGAACCTAGT (*QKI*), UCUGGACACCCUUGUUGAAUCUAUU (*BCL10*), GAAGUAGGAGAGUACUUGAAGAUGU (*CYLD*), and AGACGAAGGUUCAACAUCA (*TFEB*).

pFRT/FLAG/HA-DEST QKI (#19891) was purchased from Addgene and was subsequently cloned into a pCDH1-MSCV-EF1α-GreenPuro vector (SBI). Lentiviral GFP-expressing GIPZ sh*MALT1* (V2LHS_84221: TATAATAACCCATATACTC and V3LHS378343:

TCTTCTGCAACTTCATCCA) or non-silencing short hairpin control (shc) were purchased from Open Biosystems. Lentiviral particles were obtained from pSPAX2 and pVSVg co-transfected HEK-293T cells and infected as previously described^55^. pFRT/FLAG/HA-DEST QKI was a gift from Thomas Tuschl (Addgene plasmid #19891)^56^, pRluc-LC3wt and pRluc-LC3BG120A were a gift from Marja Jaattela (Addgene plasmid #105002 and #105003)^57^. They were introduced in GSCs using Neon electroporation system (MPK5000) according to manufacturer’s instructions (Life Technologies).

### Antibodies and Reagents

Cathepsin Inhibitor 1, Rapamycin, and Mepazine were purchased from Selleckchem, Tocris Bioscience, and Chembridge, respectively. Bafilomycin A1, Cycloheximide, phorbol myristate acetate (PMA), Pepstatin A, fluphenazine, cyamemazine, chlorpromazine, pipotiazine, alimemazine, promethazine, and doxylamine were all from Sigma-Aldrich. zVRPR.fmk was purchased from Enzo Life Sciences. Q-VD-OPh and Tumor Necrosis Factor-alpha (TNFα) were obtained from R&D Systems. Ionomycin was purchased from Calbiochem. The following primary antibodies were used: NESTIN (Millipore MAB5326), SOX2 (Millipore AB5603), GAPDH (Santa Cruz SC-25778 and SC-32233), TUBULIN (Santa Cruz SC-8035), MALT1 (Santa Cruz SC-46677), LAMP2 (Santa Cruz SC-18822), BCL10 (Santa Cruz SC-13153), BCL10 (Santa Cruz SC-5273), CYLD (Santa Cruz SC-137139), HOIL1 (Santa Cruz SC-393754), QKI (Santa Cruz SC-517305), PARP (Santa Cruz SC-8007), IKB α (CST 9242), p-S32/S36-IKB α (CST 9246), P62 (CST 5114), mTOR (CST 2983), p-S473-AKT (CST 4060), AKT (CST 9272), p-S235/S236-S6 (CST 2211), p-T183/Y185-JNK (CST 9255), JNK (CST 9258), p-S757-ULK1 (CST 6888), LC3B (CST 3868), p-T37/T46-4E-BP1 (CST 2855), p-T70-4E-BP1 (CST 9455), p-S65-4E-BP1 (CST 9451), 4E-BP1 (CST 9644), eIF4E (CST 2067), TOM20 (CST 42406), p-T421/S424-p70S6K (CST 9204), p70S6K (CST 14130), EEA1 (BD Bioscience 610456), CSTD (BD Bioscience 610800), PEX1 (BD Bioscience 611719), PECAM (BD Bioscience 557355), TFEB (Bethyl A303-673A), PDI (Abcam ab2792), GM130 (Abcam ab52649), QKI (Atlas HPA019123), CSTD (Atlas HPA063001), and FLAG (Sigma F1804). HRP-conjugated secondary antibodies (anti-rabbit, mouse Ig, mouse IgG1, mouse IgG2a, and mouse IgG2b) were purchased from Southern Biotech. Alexa-conjugated secondary antibodies were from Life Technologies.

### Tumorsphere Formation

To analyze tumorsphere formation, GSCs (100/µL) were seeded in triplicate in NS34 media as previously described^15^. Cells were dissociated manually each day to reduce aggregation influence, and maintained at 37°C 5% CO_2_ until day 5 (day 4 for siRNA). Tumorspheres per field of view (fov) were calculated by counting the total number of tumorspheres in 5 random fov for each well. The mean of each condition was obtained from the triplicates of 3 independent experiments.

### Limiting Dilution Assays

In order to evaluate the self-renewal of GSCs, limited dilution assays (LDA) were performed as previously described^58^. GSCs were plated in a 96-well plate using serial dilution ranging from 2000-1 cell/well with 8 replicates per dilution and treated as indicated. After 14 days, each well was binary evaluated for tumoursphere formation. Stemness frequency was then calculated using ELDA software^59^. The mean stemness frequency for each treatment was calculated by averaging across 2 independent experiments.

### Cell Viability

Cell viability of GSCs #1, #4, #9, and #12 was evaluated through the Uptiblue reagent (Interchim), a colormetric growth indicator based on detection of metabolic activity. Cells were plated at 5000 cells per well in triplicate with vehicle (DMSO) or MPZ. 24 hours later, Uptiblue was added to each well at a concentration of 10% v/v. Cells were maintained at 37°C 5% CO_2_ until analysis at 48 hours post-treatment. Absorbance was measured at 570 and 600 nm on a FluStarOptima (BMG Labtech) plate reader, and the percentage of cell viability calculated according to the manufacturers’ instructions. Cell viability was also measured using Cell Titer-Glo luminescent cell viability assay (Promega), according to the manufacturers’ protocol. Briefly, cells were seeded at 5000 cells per well in triplicate indicated treatment. Two days later, 100 µL of Cell Titer-Glo reagent was added to each condition, cells were shaken vigorously, using an orbital shaker, to aid in their lysis, and then luminescence was measured on a FluStarOptima (BMG Labtech) plate reader. For differentiated GSC, DGC#1 and GSC#4, cell survival was measured using the colormetric MTT assay (1-(4,5-dimethylthiazol-2-yl)-3,5-diphenylformazan, thiazolyl blue formazan, Sigma), as previously described^15^.

### ELISA

10.10^6^ GSCs were cultured with 20 µM MPZ or DMSO and culture media was collected at 8 hours, centrifuged and filtered. Human CTSD ELISA (Millipore #1317) was performed on the culture media according to the manufacturer’s instructions.

### Animal Procedures

Tumor inoculation was performed on female Balb/C nude mice (Janvier) aged 6-7 weeks, as described previously^10^. Animals were randomly assigned to each group and group housed in specific pathogen free (SPF) conditions at 24°C on a 12-hour day-night cycle. At all times, animals were allowed access to standard rodent pellets and water *ad libitum*. Mice were subcutaneously injected in each flank with 10^6^ GSC#9 in 100 µL of PBS and growth factor-free matrigel (Corning). Once tumors were palpable, mice were injected intraperitoneally daily with MPZ (8 mg/kg) or vehicle (DMSO) for two consecutive weeks. Tumor size was measured daily during treatment and for one week following treatment withdrawal, with calipers and tumor volume calculated using the following equation (width^2^×length)/2.

### Luciferase assays

Rluc-LC3B Luciferase assay was performed as previously described^57^. Briefly, GSC#9 was transfected with 1 μg plasmid using a Neon Transfection System (Life Technologies). 24 hours later, cells were treated for 4 hours with DMSO or MPZ and then assayed using Dual-Glo Luciferase assay system according to the manufacturers’ guidelines (Promega). Luminescence was measured on a FluStarOptima (BMG Labtech) plate reader.

### Flow Cytometry

For EdU analysis, cells we incubated with EdU (10 µM) for 2 hours followed by fixation and Click-it reaction according to the manufacturers’ protocol (#C10424 Life Technologies). For Propidium Iodide (PI) staining, cells were incubated for 15 minutes at room temperature with PI (100 μg/mL, V13242 from Life Technologies) following treatment with DMSO or MPZ (20 μM) according to manufacturer’s protocol. For pS473-AKT analysis, 10^6^ cells/condition were treated for 1 hour with 20 µM MPZ or DMSO. Cells were fixed with BD Phosflow fixation buffer for 10 minutes at 37°C (#558049 BD Bioscience). Cells were then permeabilized for 15 minutes using BD Phosflow perm buffer at 4°C. Cells were incubated for 1 hour with antibody (PE-p-S473-AKT #560378 BD Biosciences). Flow Cytometry analyses were performed on FACsCalibur (BD Biosciences, Cytocell, SFR Francois Bonamy, Nantes, France) and processed using FlowJo software (BD Biosciences).

### Immunostaining

After treatment, cells were seeded onto poly-lysine slides, fixed for 10 min with 4% PFA diluted in PBS, permeabilized in 0.04% Triton X100 and blocked with PBS-BSA 4% prior to 1 hour primary antibody incubation. After PBS washes, cells were incubated with AlexaFluor-conjugated secondary antibodies for 30 minutes (Life Technologies). Next, cells were incubated with DAPI for 10 minutes and mounted with prolong gold anti-fade mounting medium (Life Technologies). For Lysotracker Red DND-99 staining cells were incubated with 50 nM Lysotracker during the last 30 minutes of treatment, and cells were fixed for 10 minutes in 4% PFA. To monitor changes in lysosomal enzyme activity, DQ-Ovalbumin assay was performed, as previously described^60^. Cells were incubated with 10 µg/mL DQ-Ovalbumin for 1 hour at the end of treatment. Cells were then fixed for 10 minutes in 4% PFA. For transferrin uptake assay, following treatment, cells were washed in medium, and incubated with Alexa596-conjugated transferrin (25 µg/mL, Life Technologies) for 30 minutes at 37°C. Cells were then acid washed for 40 seconds and fixed for 10 minutes in 4% PFA. Mouse tissue sections, 7 μm of thickness, were obtained after cryosectioning of xenograft tumor embedded in OCT (Leica cryostat, SC3M facility, SFR Francois Bonamy, Nantes, France). Mouse tissue sections and human GBM samples from patients (IRCNA tumor library IRCNA, CHU Nantes, Integrated Center for Oncology, ICO, St. Herblain, France) were stained as followed. Sections were fixed 20 min in 4% PFA, permeabilized 10 min with PBS-Triton 0.2% and blocked with 4% PBS-BSA 2 hours prior to staining. Primary antibodies were incubated overnight at 4°C. All images were acquired on confocal Nikon A1 Rsi, using a 60x oil-immersion lens (Nikon Excellence Center, Micropicell, SFR Francois Bonamy, Nantes, France). Structure illumination microscopy (SIM) images were acquired with a Nikon N-SIM microscope. Z-stacks of 0.12 μm were performed using a 100x oil-immersion lens with a 1.49 aperture and reconstructed in 3D using the NIS-Element Software. All images were analyzed and quantified using the Image J software.

### Immunoblotting and Immunoprecipitation

Cells were harvested with cold PBS followed by cellular lysis in TNT lysis Buffer (50 mM TRIS pH7.4, 150 mM NaCl, 1% Triton X-100, 1% Igepal, 2 mM EDTA, supplemented with Protease Inhibitor (Life Technologies)) for 30 minutes on ice. Samples were centrifuged at 8000g to remove insoluble fraction. Tissue samples were lysed in RIPA lysis buffer for 2 hours under agitation, following homogenization with mortar and pestal. Lysates were cleared in centrifuge at max speed for 30 minutes. Cytosol and nuclei separation were performed as previously described^61^. Briefly, cells were lysed in Buffer A (HEPES 10 mM, KCl 10 mM, EDTA 0.1 mM, EGTA 0.1 mM, DTT 1 mM, Na_3_VO_4_ 1 mM, plus protease inhibitor) on ice for 5 minutes and then Buffer A + 10% Igepal was added for 5 minutes. Samples were centrifuged at 1000g for 3 minutes. Soluble fraction was cleared at 8000g. Immunoprecipitation was performed as previously described^55,61^. Briefly, cells were lysed in TNT lysis buffer for 30 minutes and cleared by centrifugation at 8000g. Samples were precleared by a 30 minute-incubation with Protein G agarose (Sigma), and then incubated for 2 hours at 4°C with Protein G agarose and 5 µg of indicated antibodies. Protein concentrations were determined by BCA (Life Technologies). Equal amount of 5-10µg proteins were resolved by SDS-PAGE and transferred to nitrocellulose membranes (GE Healthcare). Membranes were revealed using a chemiluminescent HRP substrate (Millipore) and visualized using the Fusion imaging system (Vilber Lourmat).

### Electron Microscopy

After treatment, 1 volume of warm 2.5% glutaraldehyde (0.1M PB Buffer, pH 7.2, 37°C) was added to 1 volume of cell suspension for 5 min, RT. Fixative was removed by centrifugation, and cells were treated 2.5% glutaraldehyde for 2 hours, RT. Samples were then stored at 4°C in 1% paraformaldehyde until processed. After washes (10 min ×3), cells are post-fixed by 1% OsO4/1.5% K 3[Fe(CN) 6] for 30 min following washed by ddH2O 10 min ×3, then dehydrated by 50%, 70%, 80%, 90%, 100% ethanol, 100%ethanol/100% acetone (1:1) for 5 min, 100% acetone for 3 min. Cells were infiltrated by 100% acetone/pure resin 1:1, 1:2, 1:3 for 1h, pure resin overnight, pure resin for 1h, then cells were embedded in the pure resin and polymerized at 60°C for 48h. 70nm sections were stained by uranyl acetate and lead citrate then observed under TEM at 80kV (Technology Center for Protein Sciences, School of Life Sciences, Tsinghua University, Beijing, China).

### RNA-Seq analysis

5.10^6^ GSC#9 were treated with vehicle (DMSO) and MPZ (20 µM) for 4 hours, in three biological replicates and snap-frozen on dry ice. RNA extraction (all RIN>9.0), library preparation, RNAseq and bioinformatics analysis was performed at Active Motif (Carlsbad, California, USA). Briefly, 2 µg of total RNA were isolated using the Qiagen RNeasy Mini Kit (Qiagen) and further processed in Illumina’s TruSeq Stranded mRNA Library kit (Qiagen). Libraries are sequenced on Illumina NextSeq 500 as paired-end 42-nt reads. Sequence reads are analyzed with the STAR alignment – DESeq2 software pipeline described in the Data Explanation document. The list of differentially expressed genes from DESeq2 output were selected based on 10% adjusted P-value level and a FDR of 0.1 (please see Fig 4a, 4d). Gene ontology and KEGG pathway enrichment analysis was done using DAVID bioinformatics resources portal.

### qPCR

3.10^6^ GSC#9 were treated with vehicle (DMSO) and MPZ (20 µM) for 4 hours, in three biological replicates and were snap-frozen. RNA extraction was done using Qiagen RNeasy kit. Equal amounts of RNA were reverse transcribed using the Maxima Reverse Transcriptase kit (ThermoFisher) and 30 ng of the resulting cDNA was amplified by qPCR using PerfeCTa SYBR Green SuperMix Low ROX (QuantaBio).

The following primers were used VGF forward GACCCTCCTCTCCACCTCTC, VGF reverse ACCGGCTCTTTATGCTCAGA, GNS forward CCCATTTTGAGAGGTGCCAGT, GNS reverse TGACGTTACGGCCTTCTCCTT, HEXA forward CAACCAACACATTCTTCTCCA, HEXA reverse CGCTATCGTGACCTGCTTTT, GLA forward AGCCAGATTCCTGCATCAGTG, GLA reverse ATAACCTGCATCCTTCCAGCC, CTSD forward CAACAGCGACAAGTCCAGC, CSTD reverse CTGAATCAGCGGCACGGC, LAMP2 forward CGTTCTGGTCTGCCTAGTC, LAMP2 reverse CAGTGCCATGGTCTGAAATG, LAMP1 forward ACCTGTCGAGTGGCAACTTCA, LAMP1 reverse GGGCACAAGTGGTGGTGAG, CSTB Forward AGTGGAGAATGGCACACCCTA, CSTB reverse AAGAAGCCATTGTCACCCCA, CTSS forward GCCTGATTCTGTGGACTGG, CTSS reverse GATGTACTGGAAAGCCGTTG, LC3B forward GCTCATCAAGATAATTAGAAGGCG, LC3B reverse CTGGGAGGCATAGACCATGT, ACTB forward GGACTTCGAGCAAGAGATGG, ACTB reverse AGCACTGTGTTGGCGTACAG, HPRT1 forward TGACACTGGCAAAACAA TGCA, HPRT1 reverse GGTCCTTTTCACCAGCAAGCT. Data was analyzed using the 2-ΔΔCt methods and normalized by the housekeeping genes ACTB and HPRT1.

### Statistics

Data are representative of at least three independent experiments, unless otherwise stated. Statistical analysis was performed with GraphPad Prism5 using One-way analysis of variance (ANOVA), two-way ANOVA or an unpaired two-tailed *t*-test (Student’s t test). For each statistical test, p value of <0.05 was considered significant.

## Acknowledgements

We thank SOAP team members (Nantes, France). We thank Micropicell, Cytocell and UTE facilities (SFR Santé François Bonamy, Nantes, France).

This research was funded by Fondation pour la Recherche Medicale (Equipe labellisée DEQ20180339184), Fondation ARC contre le Cancer (JG PJA20171206146), Ligue nationale contre le cancer comités de Loire-Atlantique, Maine et Loire, Vendée, Ille-et-Vilaine (JG, NB), Région Pays de la Loire et Nantes Métropole under Connect Talent Grant (JG), and SIRIC ILIAD (INCa-DGOS-Inserm_12558). KAJ and TD both received PhD fellowships from Nantes Métropole; GAG and AT hold postdoctoral fellowships from Fondation de France and Fondation ARC, respectively.

## Author Contribution

KAJ, JG, NB, conception and design, acquisition of data, analysis and interpretation of data, drafting or revising the article; GAG, CM, YL, AT, EHW, KT, TD acquisition of data, analysis and interpretation of data; JSF conception and interpretation of data. All authors approved the manuscript. All data needed to evaluate the conclusions in the paper are present in the paper and/or the Supplementary Materials. Additional data related to this paper may be requested from the authors.

## Competing interests

The authors declare that they have no competing interests.

