## supplemental Figures S1-S4 for "Control of the Endo-Lysosome Homeostasis by the Paracaspase MALT1 regulates Glioma Cell Survival"

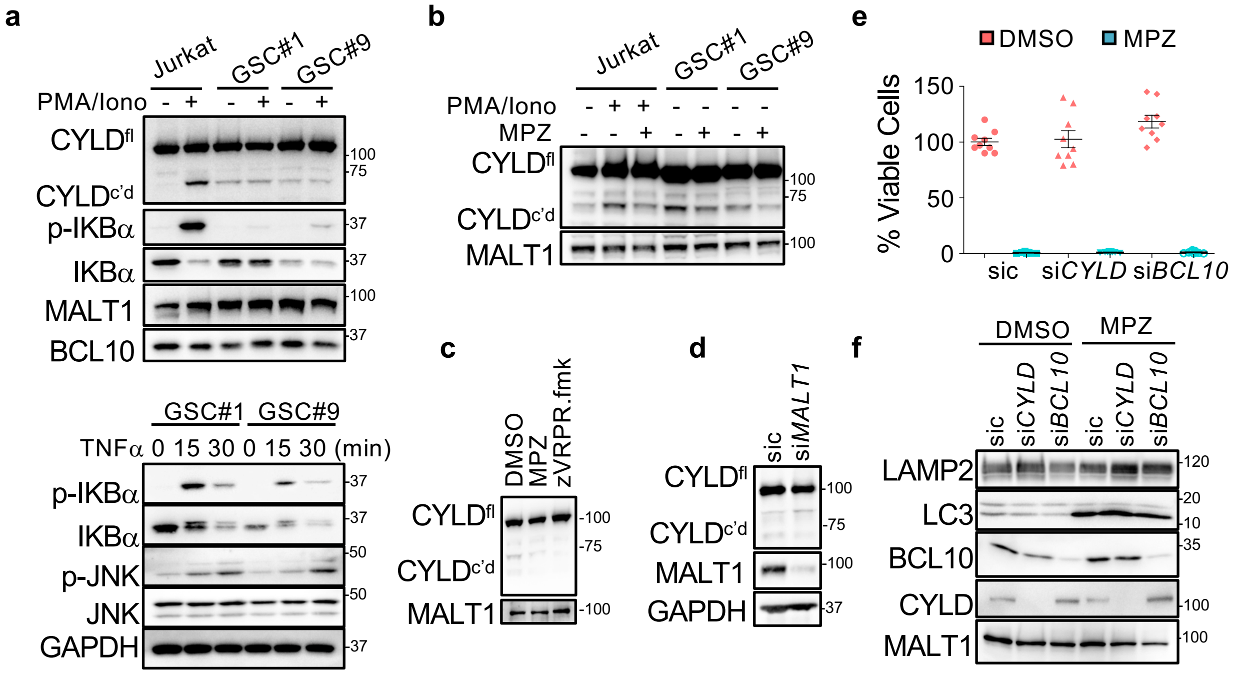


**Supplementary Figure 1. Blockade of MALT1 Activity in Glioblastoma Cells.** (a) Jurkat T-cells and GSC#1 and GSC#9 were stimulated with PMA (20 ng/mL) and Ionomycin (Iono, 300 ng/mL) for 30 minutes. Total protein lysates were analyzed by western-blot for CYLD (full length FL and cleaved c'd), p-IKBα and IKBα. MALT1 and BCL10 served as loading controls. Alternatively, total protein lysates from GSC#1 and #9 challenged with TNFα (10 ng/mL, for the indicated times) were analyzed by western-blot for p-IKBα, IKBα and p-JNK. Total JNK and GAPDH served as loading controls. (b) Jurkat T-cells and GSC#1 and GSC#9 were treated with vehicle (DMSO) and mepazine (MPZ, 20 µM) for 4 hours. PMA/Ionomycin mixture was also administered to Jurkat cells for the last 30 minutes. Total protein lysates were analyzed by western-blot for CYLD (full length FL and cleaved c'd). MALT1 served as a loading control. (c) Western-blot analysis of CYLD in total protein lysates from GSC#9 treated for 4 hours with vehicle (DMSO), MPZ (20 µM) or zVRPR.fmk (75 µM). MALT1 served as a loading control. (d) Western-blot analysis of CYLD and MALT1 in total protein lysates from GSC#9 transfected with non-silencing RNA duplexes (sic) or *MALT1* targeting duplexes (si*MALT1*). GAPDH served as a loading control. (e) Cell viability was measured using Cell TiterGlo in GSC#9 treated with vehicle (DMSO) or MPZ (20 µM) and transfected priory with sic, si*CYLD*, and si*BCL10*. Data were normalized to the vehicle-treated controls and are presented as the mean $\pm$ s.e.m of 3 independent experiments in triplicate. (f) Western-blot analysis of LAMP2, LC3B, CYLD and BCL10 in total protein lysates from GSC#9 transfected with sic, si*CYLD*, and si*BCL10* and treated 6 hours with vehicle (DMSO) or MPZ (20 µM). MALT1 served as a loading control. All data were repeated in 3 independent experiments.

**a**

| | Chemical Name (DRUG) | Structure | Cell Death | Lysosome Increase | MALT Inhibition |
|---|---|---|---|---|---|
| anti-psychotic | Mepazine (PACATAL) | 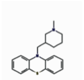 | +++ | ++ | +++ |
| | Fluphenazine (MODECATE) | 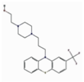 | +++ | ++ | +++ |
| | Chlorpromazine (LARGACTIL) | 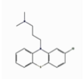 | ++++ | ++ | ++++ |
| | Cyamemazine (TERCIAN) | 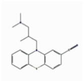 | + | ++ | ++ |
| | Pipotiazine (PIPORTIL) | 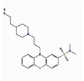 | - | + | - |
| anti-histamine | Doxylamine (DONORMYL) | 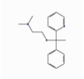 | - | N/A | - |
| | Alimemazine (THERALENE) | 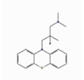 | + | + | + |
| | Promethazine (PHENERGAN) | 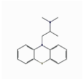 | + | ++ | + |

**b**

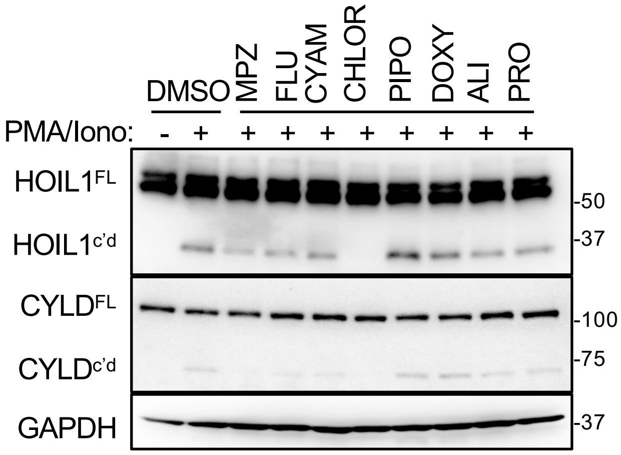


**c**

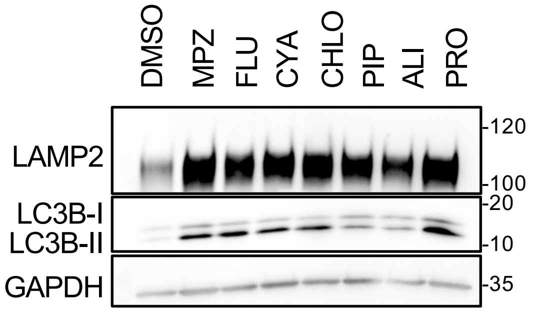


**Supplementary Figure 2. Impact of Phenothiazines on MALT1 protease activity and Lysosomes.** (a) Table summarizing eight phenothiazines used in clinics as either anti-psychotic or anti-histaminic, along with their generic and brand names (cap letters), and chemical structures. Their impact on cell death and lysosome protein increase in GSCs, and MALT1 activity in Jurkat cells is also reported. (b) Western-blot analysis of two MALT1 substrates, HOIL1 and CYLD, either full length (FL) or cleaved (c'd) in Jurkat T-cells treated with with vehicle (DMSO) or phenothiazines (20 µM CYA, CHLO, PIPO, DOXY, ALI, and PRO, 10 µM MPZ and FLU) for 30 minutes and stimulated for 30 minutes more with PMA (20 ng/mL) and Ionomycin (Iono, 300 ng/mL). GAPDH served as a loading control. (c) Western-blot analysis of LAMP2 and LC3B in equal amount of total protein lysates from GSC#9 treated for 6 hours with vehicle (DMSO) or 20 µM phenothiazines (MPZ, FLU, CYA, CHLO, ALI, PRO). GAPDH served as a loading control. All data were repeated in 3 independent experiments.

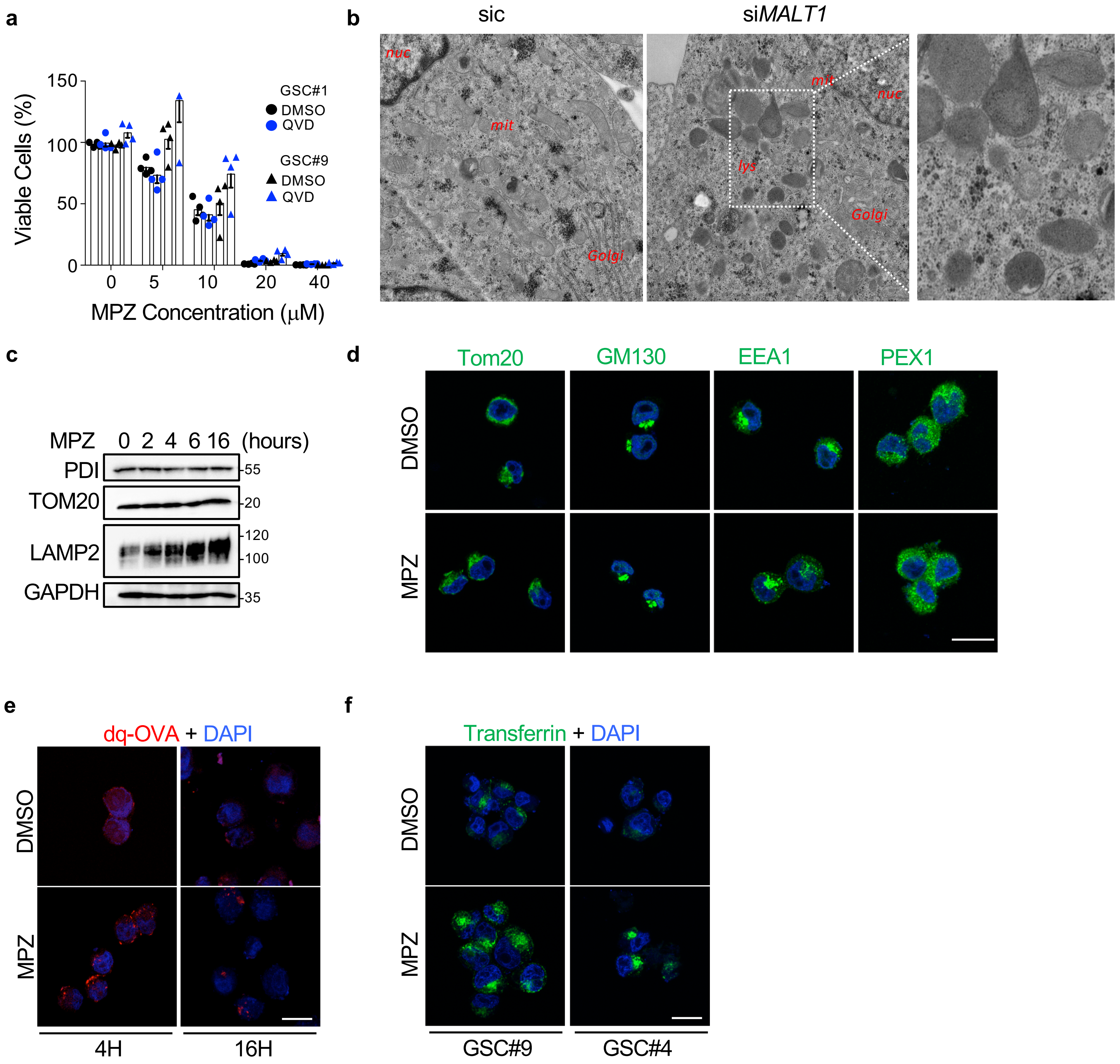


**Supplementary Figure 3. Impact of MALT1 Inhibition on intracellular Organelles.** (a) Cell viability was measuring using Cell TiterGlo in GSC#1 and GSC#9 pre-treated for 1 hour with vehicle (DMSO) or QVD (20 μM), and treated for 48 hours more with the indicated doses of MPZ. Data were normalized to the vehicle-treated controls and are presented as the mean $\pm$ s.e.m of 4 independent experiments. (b) Transmission electron microscopy images from GSC#9 transfected with non-silencing duplexes (sic) or siRNA duplexes targeting *MALT1* (si*MALT1*). Multiple images and sections from one experiment were analyzed. All data were repeated in 3 independent experiments, unless specified. (c) Western-blot analysis of PDI, TOM20, and LAMP2 in total protein lysates from GSC#9 treated vehicle (DMSO) or MPZ (20 μM) for the indicated times. GAPDH serves as a loading control. (d) Confocal analysis of TOM20, GM130, EEA1, and PEX1 immunostaining (green) in GSC#9 treated with vehicle (DMSO) or MPZ (20 μM) for 4 hours. Nuclei (DAPI) are shown in blue. Scale bars: 10 μm. (e) Confocal analysis of dq-Ovalbumin (dq-OVA, red) in GSC#9 treated with vehicle (DMSO) or MPZ (20 μM) for 4 or 16 hours. Nuclei (DAPI) are shown in blue. Scale bars: 10 μm. (f) Confocal analysis of Transferrin uptake (green) in GSC#9 and GSC#4 treated with vehicle (DMSO) or MPZ (20 μM) for 4 hours. Nuclei (DAPI) are shown in blue. Scale bars: 10 μm.

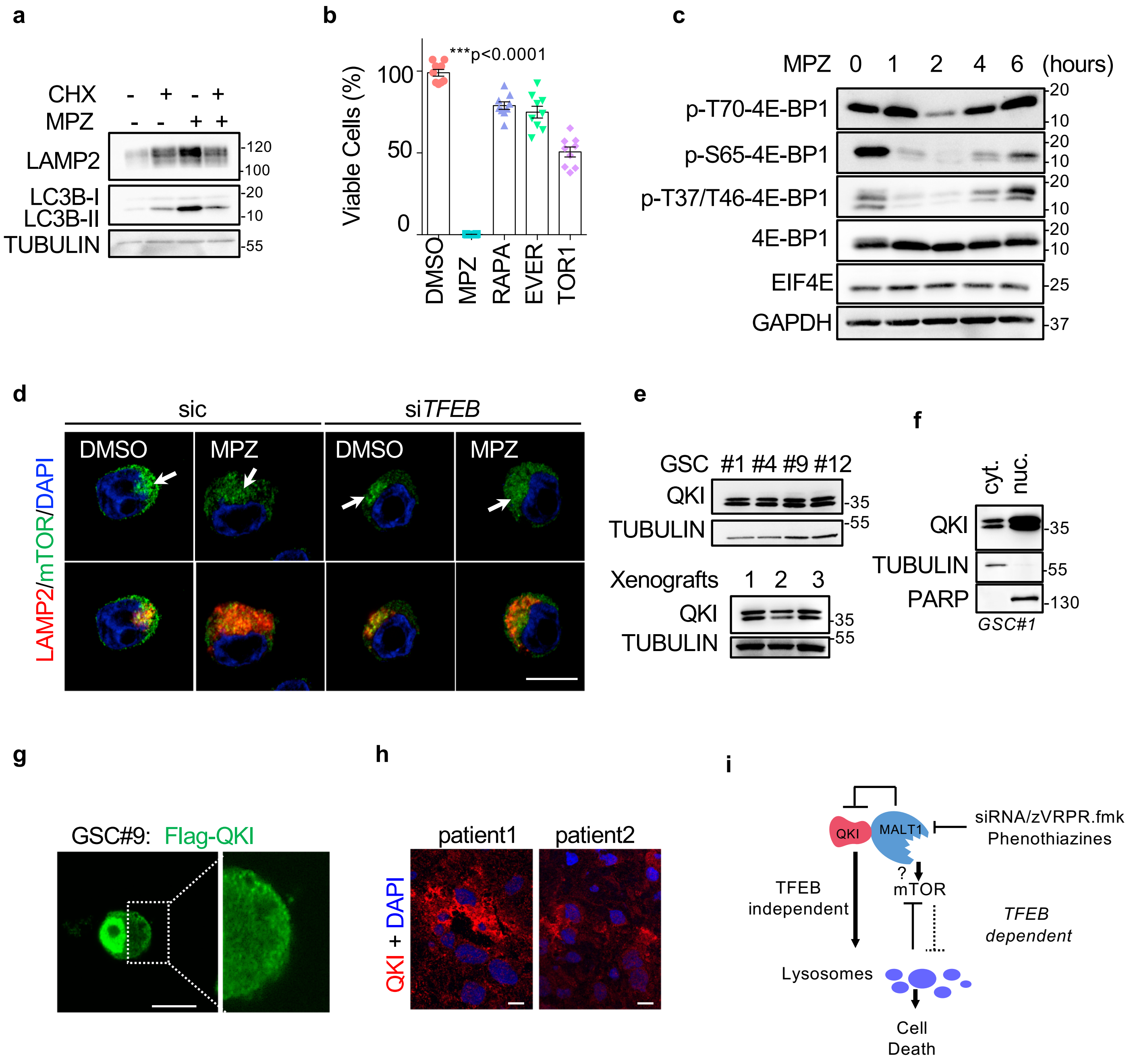


**Supplementary Figure 4. Characterization of the RNA-binding Protein QKI in Glioblastoma Cells.** (a) Western-blot analysis of LAMP2 and LC3B in total protein lysates from GSC#9 treated with vehicle (DMSO) and MPZ (20 µM) in the presence of cycloheximide (50 µg/mL), for 16 hours. TUBULIN served as a loading control. (b) Cell viability was measured using Cell TiterGlo luminescent assay in GSC#9 treated for 72 hours with vehicle (DMSO), MPZ (20 µM), Rapamycin (RAPA, 50 nM), Everolimus (EVER, 50 nM), or Torin1 (TOR1, 100 nM). Data were normalized to the vehicle-treated controls and are presented as the mean + s.e.m of 3 independent experiments in triplicate. (c) Western-blot analysis of indicated antibodies in total protein lysates from GSC #1, (d) Confocal analysis of LAMP2 (red) and mTOR (green) staining in GSC#9 transfected with sic or si*TFEB* and treated with vehicle (DMSO) or MPZ (20 µM) for 16 hours. Arrows point to LAMP2-positive area. Scale bars: 10 µm. (e) Western-blot analysis of QKI in total protein lysates from GSC #1, #4, #9, #12 and from GSC xenografted tumors. TUBULIN served as a loading control. (f) Western-blot analysis of QKI in cytosolic (cyt.) and nuclear (nuc.) cell fractionation from GSC#1. TUBULIN and PARP served as controls for each fraction. (g) Confocal analysis of Flag-QKI (green) localization in transfected GSC#9. Scale bars: 10 µm. (h) Confocal analysis of QKI immunostaining (red) in glioblastoma tissue sections from two patients. Nuclei (DAPI) are shown in blue. Scale bars: 10 µm. (i) Schematic representation of working model.
